# Co-evolution between colibactin production and resistance is linked to clonal expansions in *Escherichia coli*

**DOI:** 10.1101/2025.11.17.686825

**Authors:** Tommi Mäklin, Jessica K Calland, Sudaraka Mallawaarachchi, Harry A Thorpe, Rebecca A Gladstone, Jukka Corander

## Abstract

Specific strains of *Escherichia coli* employ the polyketide synthase island to produce a metabolite called colibactin that is implicated in colorectal tumorigenesis via its genotoxic effect on human DNA. However, the damage to host DNA is essentially collateral from ecological competition during bacterial colonisation, where colibactin is used to displace other gut bacteria. Despite intensive recent research on colibactin and its involvement in tumorigenesis, the evolutionary dynamics related to its production remain poorly characterised. We map evolution of colibactin production in *E. coli* using a large collection of high-resolution genomes and show that its introduction via a mobile element has induced multiple plasmid-mediated acquisitions of previously unrecognised colibactin immunity genes in non-producing multi-drug resistant lineages. Our study suggests that in *E. coli*, multi-drug resistance is incompatible with colibactin production due to the unbearable combined fitness costs, supported by colibactin-producing phenotypes remaining susceptible to most classes of antibiotics and thriving in regions with low antimicrobial usage. Consequently, resistant lineages circulating in these regions are under selective pressure to acquire colibactin immunity to compete with the endemic colibactin-producing bacteria during colonisation. Conversely, high antimicrobial usage selects for a colibactin susceptible antimicrobial resistant phenotype as it drives the colibactin-producing *E. coli* towards extinction. Such a duality of evolutionary strategies to become endemic in a host population structured by varying ecological pressures may hold more generally across bacterial species colonizing the human gut.

## Introduction

*Escherichia coli* with the polyketide synthase (pks) island of genes for colibactin production have recently come under intense scientific scrutiny after the discovery of a causal link between this genotoxin and mutational signatures found in colorectal cancer biopsies ^1–3^. Further research into global colorectal cancer incidence found a strong geographical correlation between the distribution of toxin-producing *E. coli* and standard of living, reporting that colibactin-producing pks positive (pks+) *E. coli* are nearly absent in low- and middle-income regions with a low incidence of colorectal cancers ^4^. This correlation is mirrored in the distribution of colibactin induced tumour signatures ^5^, conclusively showing that exposure to colibactin is a major factor in the increasing incidence of colorectal cancer in high-income regions. Colibactin may additionally contribute to the development of cancers in the urinary tract ^1,6–8^, where pks+ *E. coli* are major causes of infections ^9,10^, but the ecological and evolutionary factors driving the success and spread of pks+ *E. coli* remain unknown. Importantly, an advantage in bacterial competition at different body sites of the human host during colonisation is the main benefit from colibactin production ^11^, and it is necessary to consider the evolutionary processes taking place among key pathogenic bacteria occupying this niche to understand the natural selection acting on both production of and resistance to this toxin.

The major pks+ *E. coli* strains in high-income countries belong to the clinically relevant sequence types (ST) ST73 and ST95 that are responsible for a large number of bloodstream and urinary tract infections and neonatal meningitis cases ^9,10,12^. Somewhat puzzlingly, ST73 and ST95 are predominantly susceptible to most classes of antibiotics, despite the clear pressure to horizontally acquire resistance genes which has manifested in the recent expansions of more resistant clones such as ST69 and ST131 ^10,13,14^. This surprising pattern is likely a result of inter-strain competition between different *E. coli*, as all four STs are frequently found in asymptomatic colonization in high-income settings ^4,15^. These STs typically carry multiple plasmid associated competition mechanisms that are directly aimed at other *E. coli* ^16^ or contribute to metabolic processes ^17^ which results in differing colonization ability even in the absence of antimicrobial pressure ^18^. These factors point to more intricate evolutionary dynamics that influence the global success of pks+ and pks- *E. coli* clones.

Colibactin production in the gut is under strong regulation by environmental factors related to the availability of essential nutrients such as iron and oxygen ^11,19–21^. Being facultative anaerobes, pks+ *E. coli* employ a strategy where they switch off production of colibactin when oxygen is present ^22^ and the other fully anaerobic bacteria are unable to grow, allowing them to establish themselves in parts of the gut where oxygen is more abundant. In the more hypoxic parts, pks+ *E. coli* revert to producing colibactin to gain an edge on competing bacteria through the lethal effect of the toxin on a wide range of competing species ^23–25^.

Colibactin likely also inhibits pks- *E. coli* that lack the colibactin self-resistance gene *clbS* ^26^, corroborated by the ability of the pks+ probiotic strain Nissle 1917 to outcompete other *E. coli* ^27–29^, but conclusive proof remains elusive as the toxin is considered unstable and cannot be isolated from cultures ^30^. Nevertheless, evidence from the establishment of pks+ and pks-*E. coli* strains in the neonatal gut immediately after birth in regions with endemic pks+ *E. coli* ^4,31,32^ indirectly supports a role for colibactin in within-species competition, as co-colonization by pks+ and pks- *E. coli* is exceedingly rare ^15^. Despite this, certain pks- *E. coli* thrive in the same regions, hinting that these bacteria carry some means to either outcompete or resist the colibactin producing *E. coli*.

We present genomic and protein structural analyses that provide evidence for both chromosomal and plasmid-mediated colibactin resistance in pks- *E. coli* and elucidate the evolutionary history of colibactin production in pks+ *E. coli*. Our study is based on a longitudinally sampled representative collection of 1 991 *E. coli* isolates with high-resolution hybrid assemblies ^10,16^ that is ideal for understanding the evolutionary dynamics of the pks island. We report that pks- *E. coli* often carry a previously overlooked gene encoding a protein with highly similar structure to the colibactin self-resistance protein ClbS. The *clbS*-like gene is widespread in pandemic multi-drug resistant (MDR) lineages and has played a part in their clonal expansion within the past few decades according to its inferred date of introduction. Our results strongly suggest that co-evolution between colibactin production and resistance has contributed to shaping the global population structure of extra-intestinal pathogenic *E. coli*.

## Results

### Clinically relevant *E. coli* carry genes from two families with highly similar structure to the colibactin self-resistance protein ClbS

We first used a collection of 1 991 high-quality *E. coli* hybrid assemblies obtained from bloodstream infections in Norway between 2002–2017 (termed subsequently as the NORM collection) that were isolated without selection for any specific phenotype ^10,16^ to look for structural homologs of the colibactin self-protection protein ClbS, a part of the 19-gene polyketide synthase (pks) island that enables colibactin production in pks+ *E. coli* ^33^. We identified two families of ClbS-like proteins that bear high structural similarity to ClbS (Figure 1) and are widespread in the NORM collection (Table 1). In our collection, these genes were annotated as a DUF1706 domain-containing protein and have previously been given the name *dfsB* ^34^ (*dendritiformis* sibling bacteriocin).

**Figure 1.**
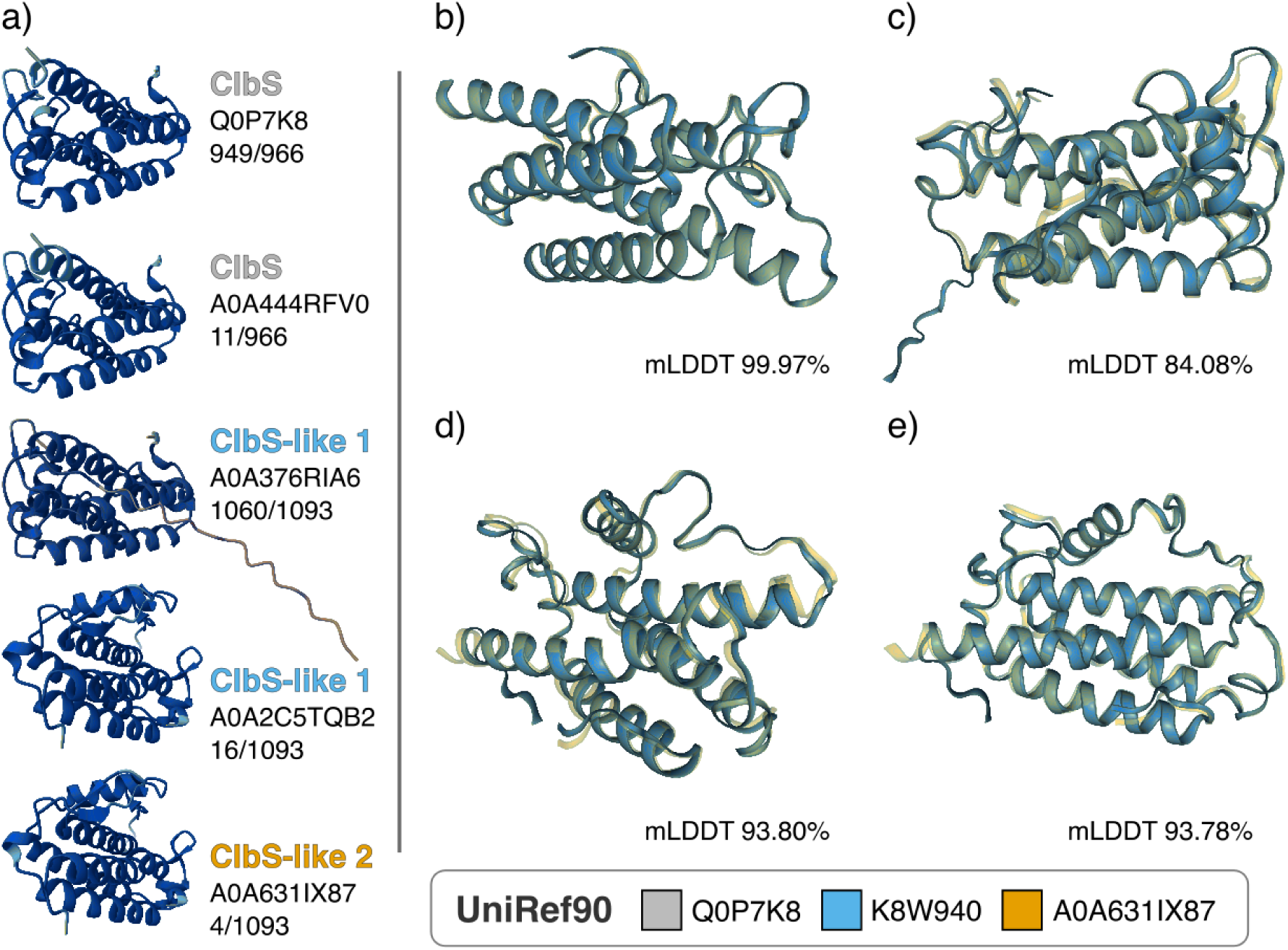
Structural comparisons between the colibactin self-resistance protein ClbS and the ClbS-like proteins present in the NORM dataset. Panel a) shows the structural predictions for the pks-island associated ClbS proteins and the two ClbS-like proteins. The colours correspond to the UniRef90 cluster for the proteins (90% sequence identity and 80% overlap). The numbers in panel a) denote the number of identifications of each ClbS or ClbS-like variant out of all ClbS or ClbS-like variants. Panels b-e show structural alignment between the ClbS protein (Q0P7K8) and the b) ClbS variant (A0A444RFV0), c) ClbS-like 1 protein (A0A376RIA6), d) ClbS-like 1 protein (A0A2C5TQB2), and e) ClbS-like 2 protein (A0A631IX87). In panels b-e, mLDDT denotes the mean local distance difference test (LDDT) values.

**Table 1.**
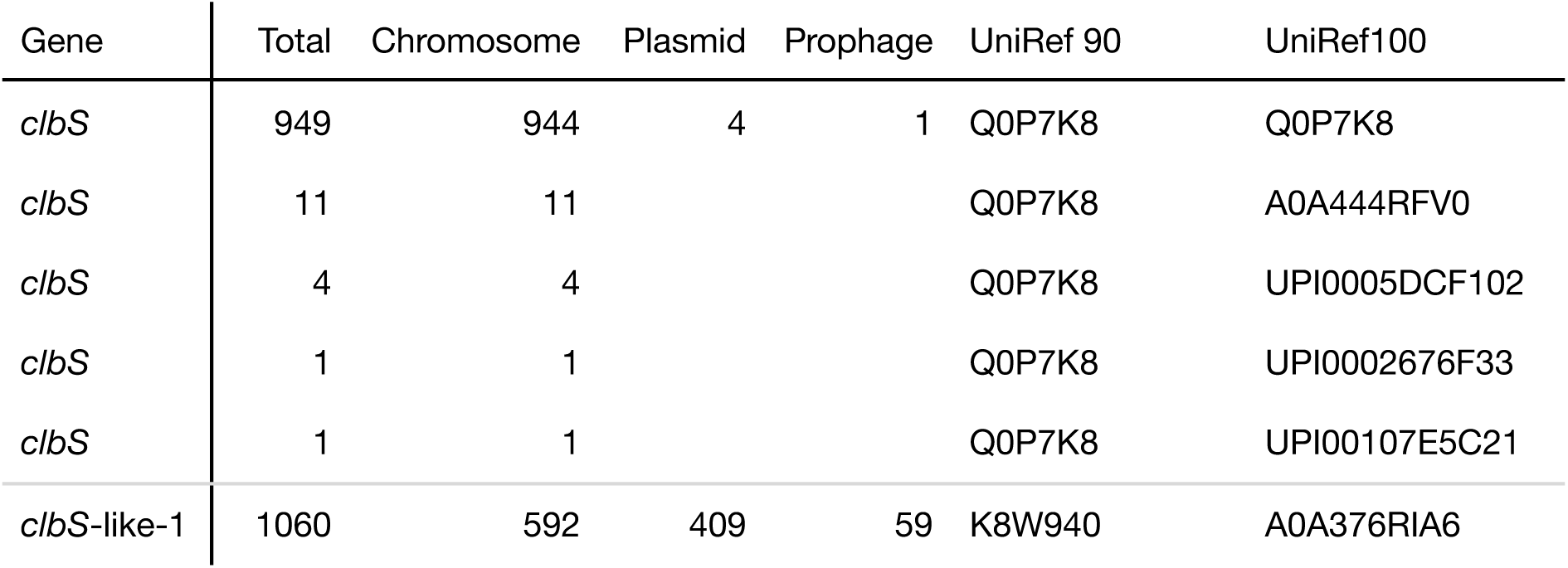
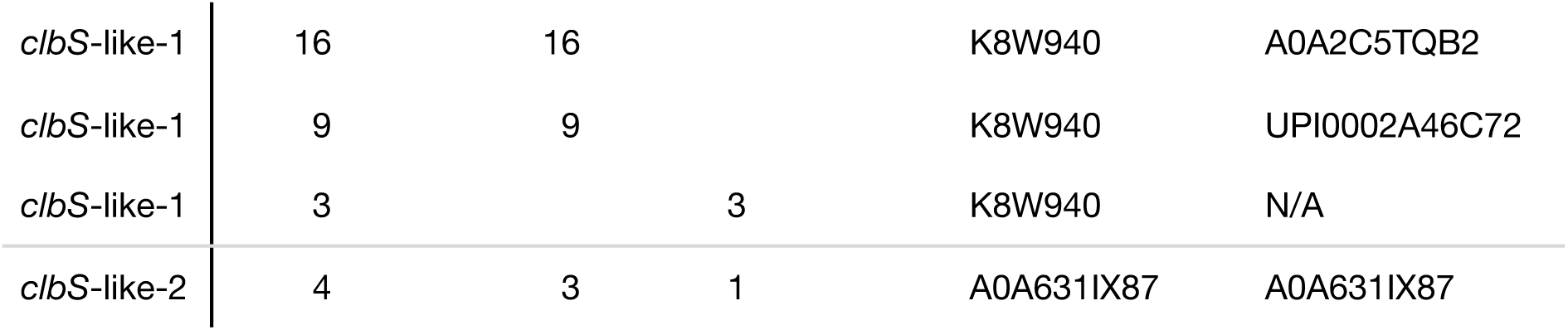
Colibactin self-resistance gene *clbS* and putative *clbS*-like gene variants from the NORM hybrid assemblies. The genes were identified either by a structural similarity or nucleotide sequence similarity. The total number of each gene, including duplicates, is indicated in the “Count” column, followed by the count of each gene in the chromosome, a plasmid, or a prophage sequence. The “UniRef” columns denote the UniRef 90%, and 100% sequence identity cluster identifiers.

The ClbS-like proteins are encoded by divergent nucleotide sequences from the *clbS* gene, with a median pairwise difference of 225 out of 522 sites (222-227 in 1%-99% percentile range with standard deviation 2.6), not including gaps, between the *clbS* and *clbS*-like genes in a multiple sequence alignment. However, the high degree of structural conservation between these structures implies that the proteins encoded by the *clbS*-like genes protect the bacterial host from colibactin and provide a fitness advantage in the presence of competing pks+ strains, whereas the divergence between nucleotide sequences of the *clbS* immunity gene present on the pks island and *clbS*-like genes suggest that these genes do not share recent evolutionary history. Furthermore, the two *clbS*-like gene families also have diverging nucleotide sequences, implying different origins and possible transmission via horizontal gene transfer. To investigate this further, we next performed a screen mapping the colibactin production genes and the putative resistance genes across the *E. coli* species phylogeny.

### Colibactin production genes are restricted to phylogroup B2 *E. coli* but *clbS*-like resistance genes transcend phylogroup boundaries via plasmids

Consistent with previous reports on pks island carriage, in NORM collection the pks island is nearly exclusive to the B2 phylogroup (Figure 2), which contains most of the major extra-intestinal pathogenic *E. coli* strains. Notably, in NORM, nearly half of the assemblies carry the pks island and it is ubiquitously present in the chromosomes of the two most common STs: ST73 and ST95 (Figure 2), while we never observed it on a plasmid. Together, the phylogeny and the exclusive chromosomal carriage of the pks island imply that the colibactin production genes were likely present in the common ancestor of the pks+ *E. coli* in phylogroup B2 and were acquired relatively far back in their evolutionary history.

**Figure 2.**
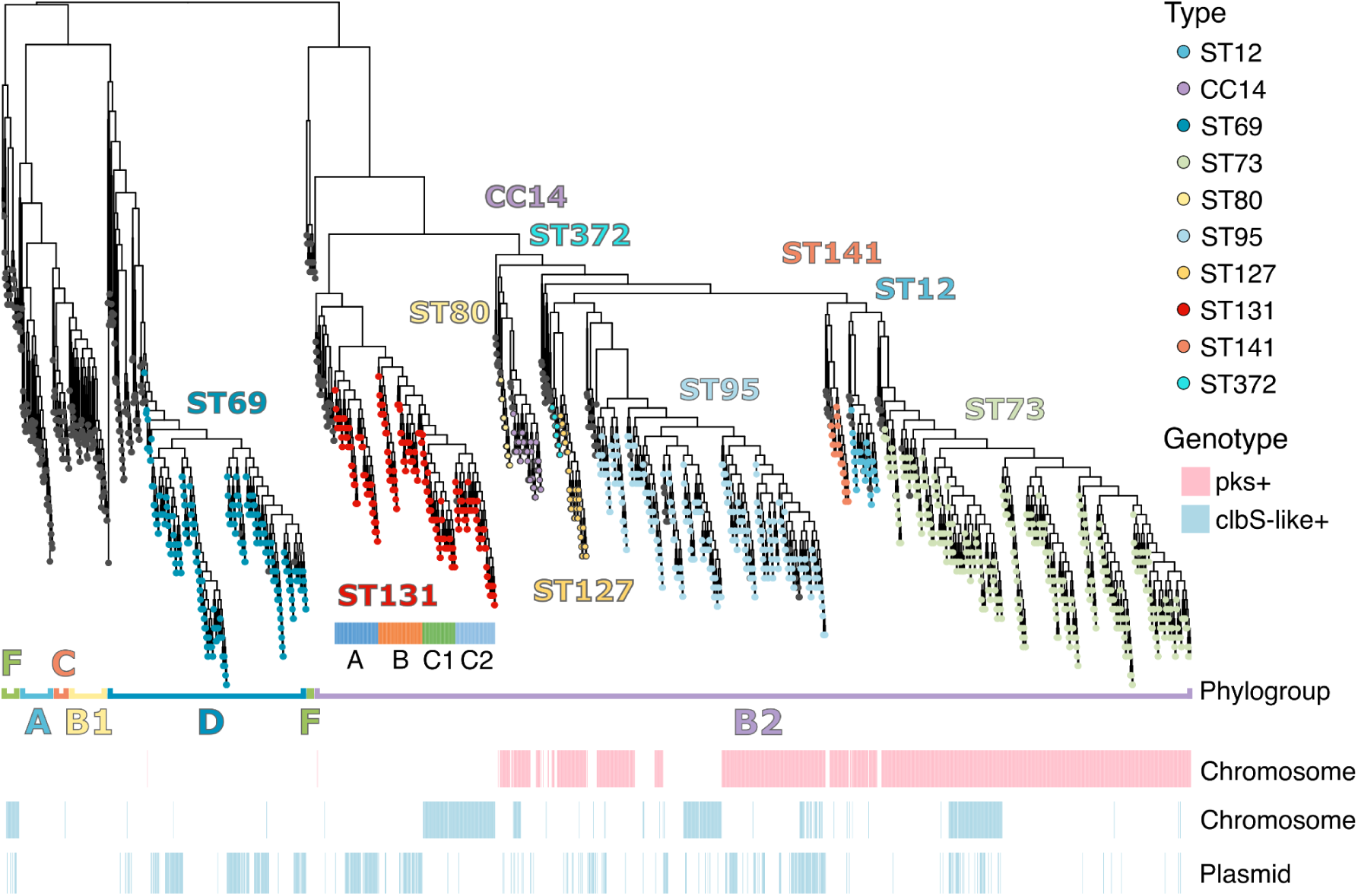
Distribution of colibactin production genes (the pks island) and putative resistance gene *clbS*-like in 1 991 NORM *Escherichia coli* isolates. Tip label colours show the multi-locus sequence type for common colibactin producing (pks+) sequence types (ST) or clonal complexes (CC). For ST131 clades A, B, C1, C2 are additionally shown. Chromosomal and plasmid carriage of pks and *clbS*-like genes are indicated below the phylogeny.

Mapping the presence of *clbS*-like genes across the same phylogeny reveals a much more dynamic pattern, with these genes both carried on plasmids and found integrated in the chromosome. In both the chromosomes and plasmids, the *clbS*-like gene is located next to a single ISL3 family transposase near otherwise unrelated genes (Supplementary Figure 1), suggesting that the chromosomal integrations are derived from a transposon that crosses phylogroup boundaries by piggybacking on mobile elements. In particular, ST69 from phylogroup D and ST131 from phylogroup B2, which are among the most successful globally disseminated multi-drug resistant (MDR) lineages ^35^, often carry the *clbS*-like genes (Figure 2). In the case of ST131, the most recently expanded clades ST131-C1 and ST131-C2 have furthermore integrated the *clbS*-like transposon in their chromosomes, showing that colibactin resistance is an essential trait for these clades. Since the primary ecological niche of these opportunistic pathogens is asymptomatic colonisation of the human gut, these strains likely frequently meet other commonly distributed *E. coli* that produce colibactin, such as ST73 and ST95. Interestingly, *clbS*-like gene carriage is also found in many pks+ isolates, and in particular in an ST95 clade that has lost the colibactin production genes (Figure 2), indicating that the previously unrecognised role of *clbS*-like genes is important across different genetic backgrounds.

### Pks island genes were present in the most recent common ancestor of pks+ lineages but colibactin resistance via a plasmid is a recent acquisition

To further investigate the evolutionary interplay between colibactin production and resistance, we looked at dated phylogenies for the common pks+ lineages in NORM (ST73, ST95 and the clonal complex (CC) 14; Figure 3) and the two most common pks-lineages (ST69, ST131; Figure 4). Furthermore, for isolates that carried the *clbS*-like gene on a plasmid, we extracted the plasmid types from a previous study ^16^ (summarised in Methods). These data confirm that the pks+ phenotype was already present in the most recent common ancestor of the pks+ ST73, ST95 and CC14 (Figure 3) but resistance without production is a more recent acquisition (Figure 4). Additionally, all five lineages carry pUTI89-like plasmids (denoted by types 2-1, 2-2, and 2-3) which contain the *clbS*-like gene but have only acquired these plasmids evolutionarily recently (towards the latter half of the 20th century). In all but two isolates, the assemblies contain only one distinct *clbS*-like plasmid. The two exceptions with multiple distinct plasmids were both from ST95 and carried a type 2-3 plasmid and an additional plasmid with unknown type, both containing a *clbS*-like gene.

**Figure 3.**
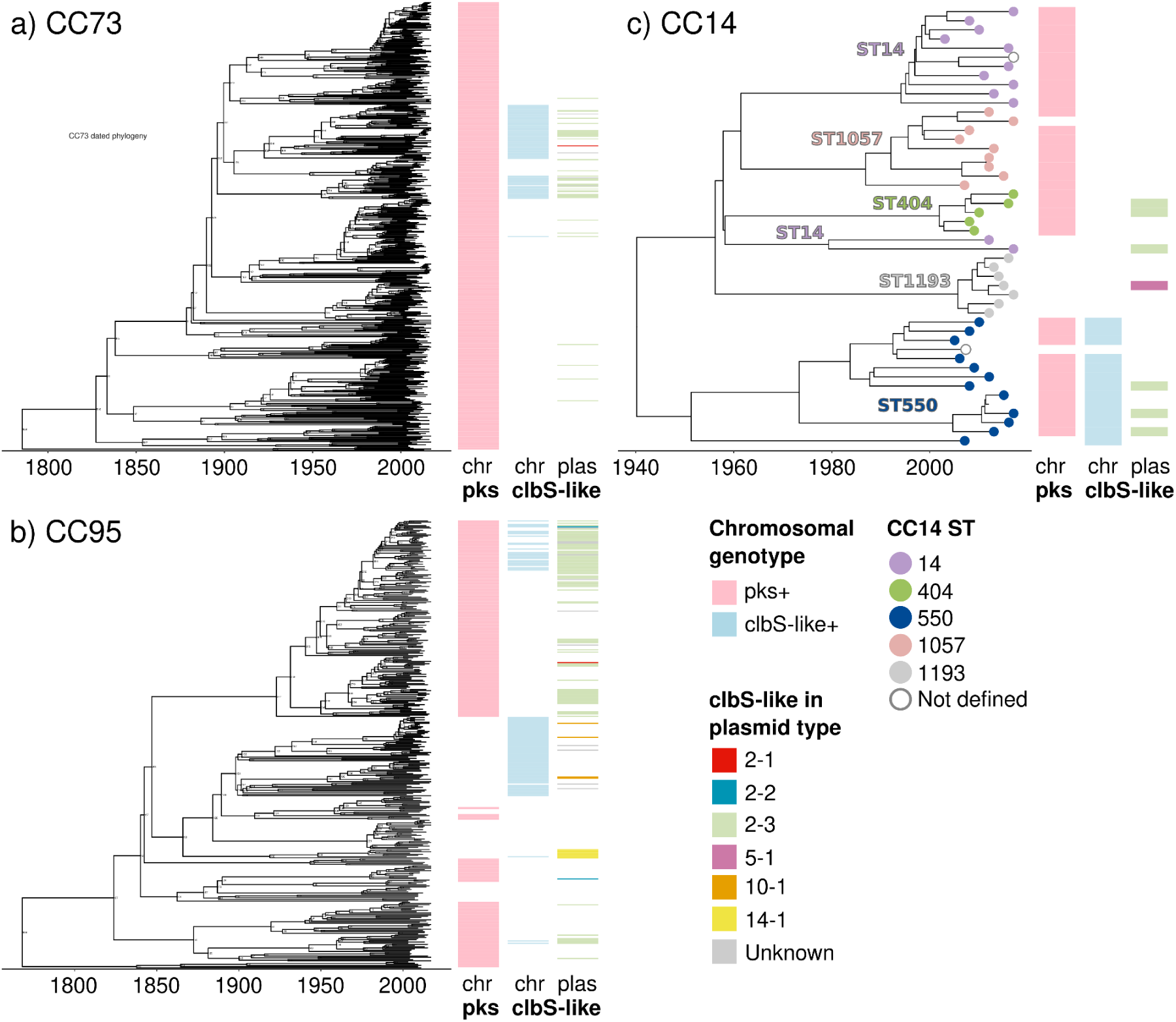
Dated phylogenies for the common pks+ sequence types in NORM showing the presence and absence of the pks island and *clbS*-like genes. Panels a-c show dated phylogenies for clonal complexes a) CC73, b) CC95, c) CC14. The horizontal axis displays the dating of each data point in years common era. The two leftmost metadata blocks show the presence of the pks island genes and the *clbS*-like gene in the chromosome. The third metadata block shows the presence of the *clbS*-like gene in a plasmid, coloured by the plasmid type (Methods). In panel c), the sequence type within the clonal complex is annotated on the tree.

**Figure 4.**
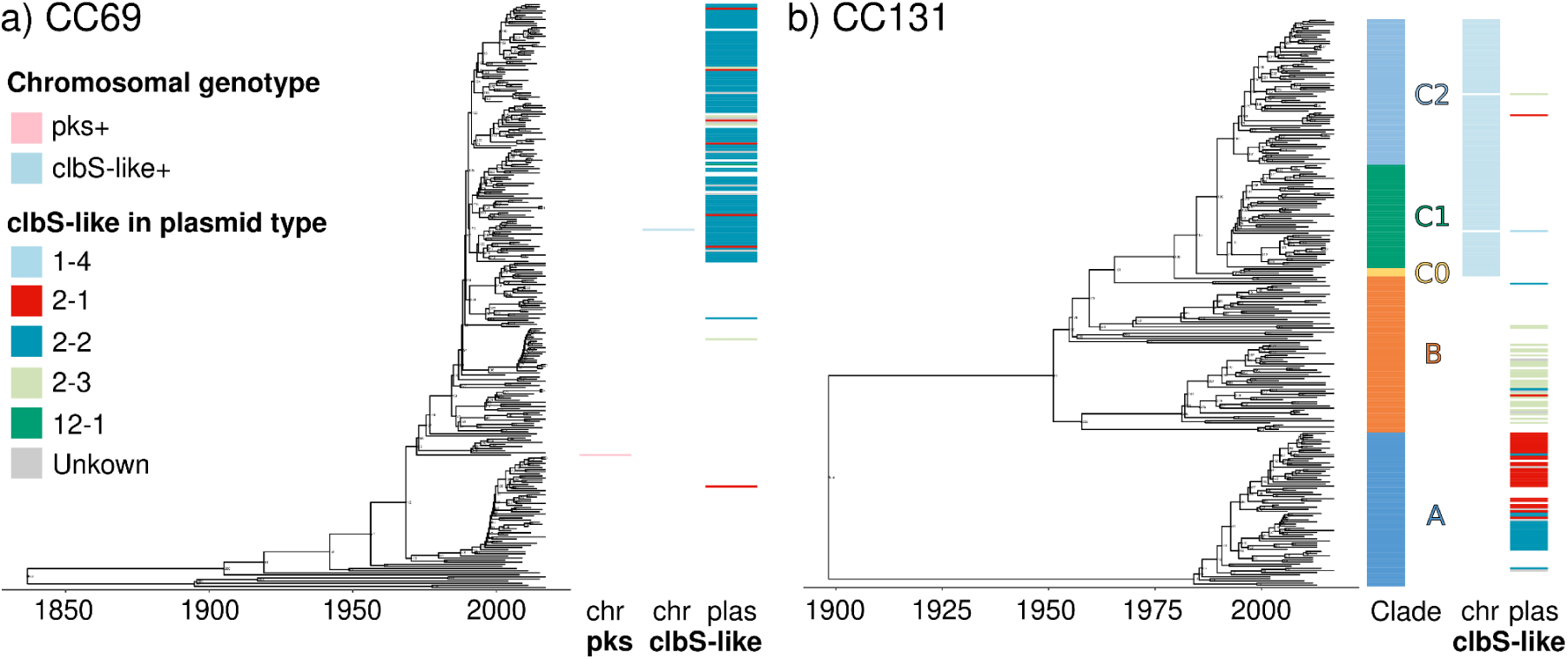
Dated phylogenies for the two most common pks- sequence types in NORM showing the presence and absence of the pks island and *clbS*-like genes. Panel a) shows a dated phylogeny for the clonal complex (CC) centered on ST69. Panel b) shows a dated phylogeny for CC131. The horizontal axis displays the dating of each data point in years common era. In Panel b), the leftmost metadata block displays the ST131 clade. The next metadata blocks show the presence of the pks island genes and the *clbS*-like gene in the chromosome or in a plasmid, coloured by the plasmid type. Plasmids with the same type that did not carry a *clbS*-like gene are not shown.

Phylogenetic analysis revealed a loss of the typically conserved pks+ phenotype in a subclade of ST95 that has integrated the *clbS*-like gene in its chromosome (Figure 3b). Based on the timing of expansion of the *clbS*-like carrying pks- clade in ST95, the pressure to lose the pks island and to in tandem acquire colibactin resistance temporally coincides with the introduction of 3rd generation of cephalosporins into widespread clinical use. This indicates a switch from a colibactin producing phenotype to resistant non-producer in a lineage which has remained a dominant cause of *E. coli* bacteria for at least several decades ^16^.

Intrigued by this observation, we looked further into the dated phylogeny for CC14 that is a notably diverse colibactin-producing clonal complex which also contains the pks- ST1193, an emerging pandemic *E. coli* clone with a similar clinical profile to ST131 C1/C2. Although the NORM collection mostly predates the recent expansion of ST1193, which started in the mid-2010’s ^36^, the collection contains 7 hybrid assemblies from ST1193. Dating the divergence of these assemblies from CC14 shows that ST1193 has recently (Figure 3c) lost the pks island genes present in all other CC14 sequence types, but has not yet acquired colibactin resistance genes. Considering the recent expansion of pks- lineages with the *clbS*-like gene within ST69, ST95, and ST131 (Figures 3b and Figure 4b), the divergence of ST1193 presents evidence for epidemiologically successful evolutionary strategies that trade colibactin production for colibactin protection genes (ST95, ST131) and multi-drug resistance (ST69, ST131, and ST1193) by acquiring the appropriate plasmids.

### The pks island is located in a chromosomal region with a high number of transposable elements, virulence determinants, and metabolism genes

Next, we looked into the chromosomal region containing the pks island genes in the pks+ *E. coli* via a pangenome graph (Figure 5). This graph shows that the colibactin production genes are always carried together with the yersiniabactin biosynthesis genes (constituting “the pks neighbourhood”) which have a dual function in iron and copper binding ^37–39^ and are located near multiple key genes related to regulation, surface proteins, and metabolic genes. In particular, the transcriptional regulators at the beginning of the pks island (*clbR* and *clbA*) are followed by the *cobU* gene, which encodes a protein involved in cobinamide salvaging and de novo adenosylcobalamin (coenzyme B_12_) synthesis ^40,41^, and *fepA*, a TonB dependent receptor for ferric enterochelin and colicin B ^42^. On the yersiniabactin side, the neighbourhood ends in the DgsA anti-repressor *mtfA* that is followed by a prophage integration site and genes related to zinc metabolism and flagellar assembly. Between the yersiniabactin and colibactin genes, pks+ *E. coli* carry multiple invasins and adhesins as well as efflux and regulation genes.

**Figure 5.**
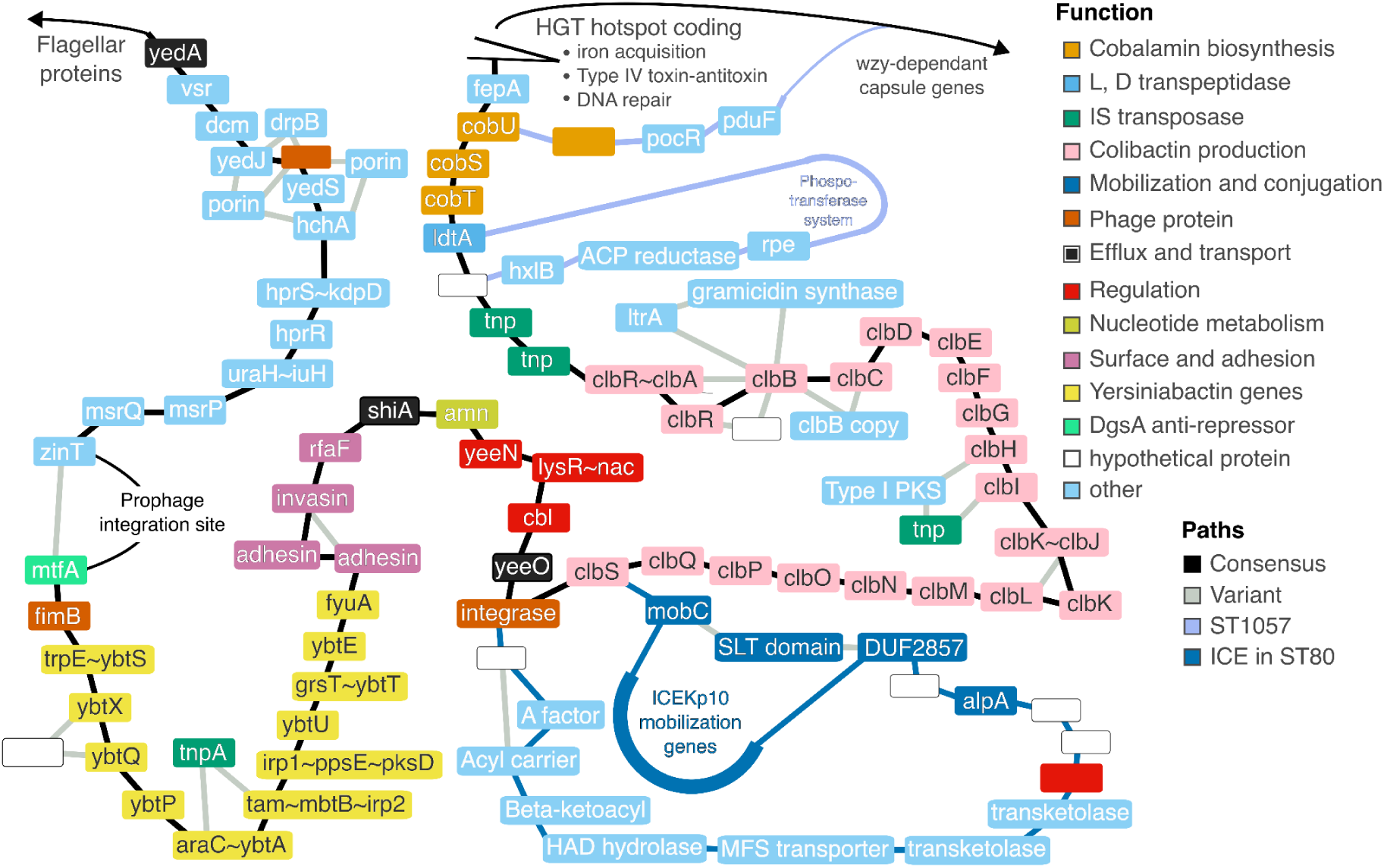
Simplified pangenome graph for the NORM hybrid assemblies showing the chromosomal neighbourhood around the pks island. The graph highlights the most common path through the region taken by the pks+ isolates (labelled “Consensus”) and the possible variants (“Variant”) . Genes in the region are coloured according to a functional grouping based on inspecting the annotations. The graph was simplified by removing nodes that are not located within the pks island or only had a single outgoing edge, and by collapsing branches containing the ICEKp10 mobilization genes, a phosphotransferase system, and a prophage integration site.

The graph also shows that apart from two branching paths present only in ST1057, a phospotransferase system between *ldtA* and a hypothetical protein and a propanediol utilization system skipping the horizontal gene transfer (HGT) hotspot after *cobU* and *fepA* (available for viewing in an interactive version of the plot provided as Supplementary Data), the configuration of the pks neighbourhood genes is typically highly conserved (Figure 5).

Contrasting this conservation, the neighbourhood itself is located between two highly variable regions, containing the phage integration site before the yersiniabactin genes and a HGT hotspot after the cob operon and *fepA*. Hence, the lack of substantial structural variation within the pks island and neighbourhood indicates that the unit as a whole is under strong purifying selection and consequently most structural changes introduced by HGT would be purged from the population.

We additionally found that the pks neighbourhood in a single CC14 assembly from ST80 was entirely derived from an integrative conjugative element (ICEKp10) from *Klebsiella pneumoniae*, that is known to carry the colibactin and yersiniabactin genes ^43^, with the mobilization machinery completely intact. Remnants of the mobilization machinery are also found in multiple ST73 assemblies, which retain the region between *mobC* and the integrase following *yeeO*. This together with the location of the neighbourhood in a variable chromosomal region suggests that the yersiniabactin and colibactin genes were likely acquired via HGT from *Klebsiella* through an ICE similar to ICEKp10.

### Probable conjugative element origin for colibactin production in *E. coli* from *Klebsiella*

We further examined the possibility of mobile element origin for colibactin production in *E. coli* by extracting the chromosomal neighbourhood for the pks island (genes from *cobU* to *mtfA*; Figure 6) from representative genomes for each sequence type in NORM and compared their contents to ICEKp10. Gene synteny (Figure 6) shows a high degree of conservation between ICEKp10 and the pks neighbourhood of an ST80 assembly, which we noted earlier contains the intact mobilization and conjugation machinery of ICEKp10.

**Figure 6.**
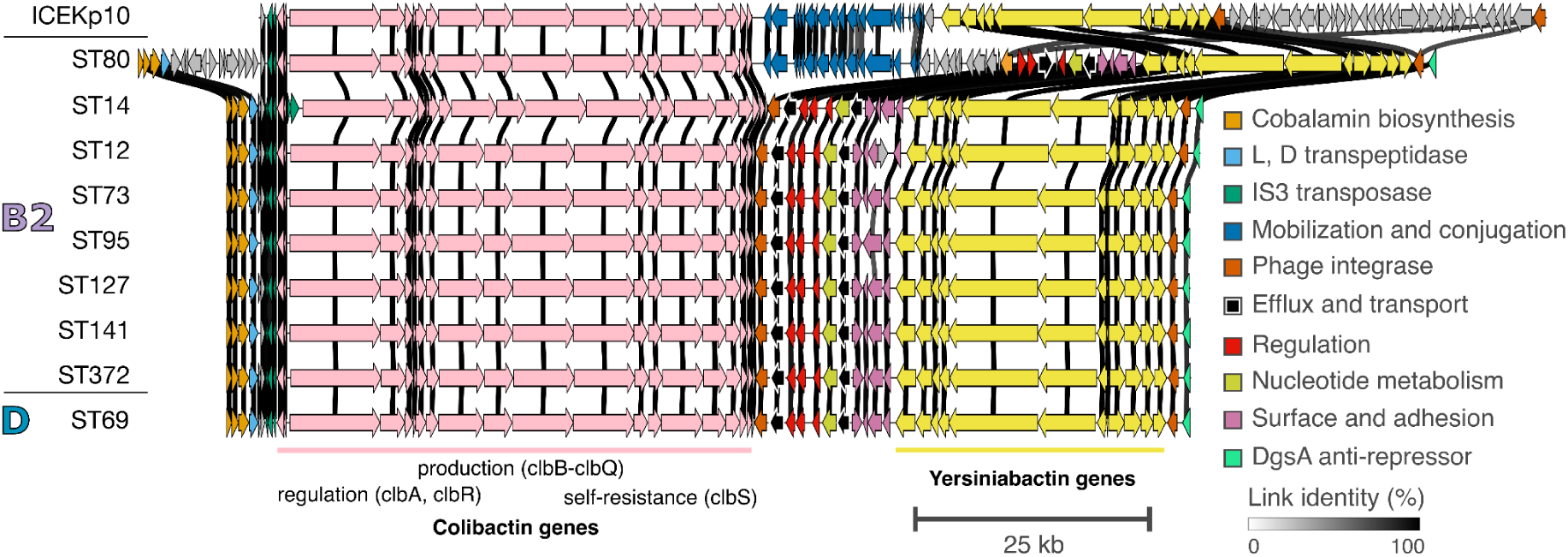
Spatial structure of the pks neighbourhood in representative *E. coli* genomes and the ICEKp10 sequence. The pks neighbourhood is defined as the region between the genes *cobU* and *mtfA*. The rows show the pks neighbourhood contents for a randomly chosen representative from each pks+ PopPUNK cluster for the NORM hybrid assemblies. Phylogroup and sequence type (ST) are indicated in the row labels. The colours correspond to a potential functional grouping of the genes, performed manually based on the genes present. Genes that were present in only a single representative or could not be assigned to a group are coloured gray. The horizontal scale bar denotes a 25 000 nucleotides long sequence. The grayscale shading of the links shows sequence identity between linked sequences, with links that had less than 70% identity in all occurrences hidden.

Considering that ST80 is closely related to clonal complex 14 (Figure 2), which our phylogenetic analysis indicates as a possible donor lineage of the pks island genes in phylogroup B2 *E. coli* (Figure 2), this means that the pks island genes in *E. coli* very likely originate from an ICE element that is amenable to interspecies jumps involving *Klebsiella*.

In the other pks+ *E. coli* lineages, the structure of the pks neighbourhood is largely conserved, with the main gene clusters almost exactly preserved among all of them (Figure 6). Somewhat surprisingly, we also found the pks island completely intact in a single phylogroup D ST69 assembly (Figure 4a). Although the pks island is not widely spread outside of phylogroup B2, the presence of the pks island in ST69 demonstrates that it can cross phylogroup boundaries in the presence of some, yet unknown, conditions. Given the apparently high fitness cost arising from integration of the pks island in most genetic backgrounds, selection for colibactin production needs to overcome this for stable carriage and further dissemination of the island. Of note, the pks+ isolate of ST69 does not belong to the pandemic MDR clade of this lineage that started expanding around 1990 ^16^ (Figure 4a), which is in congruence with the incompatibility of MDR and colibactin-producing phenotypes. Phylogenetic distribution of the pks island and the high level of structural congruence suggest a transfer happening from phylogroup B2 to D through an integrative and conjugative element.

### The pks island genes carried by *E. coli* split into two distinct and conserved majority-minority variants defined by a fusion of the *clbK* and *clbJ* genes

Lastly, we looked for variation in the sequence contents of the pks island genes (*clbA-clbS*) by performing an amino acid multiple sequence alignment on the gene sequences. Our results confirm an earlier report ^33^ which found that the only major variation in the pks island genes is a fusion of two large genes from the non-ribosomal peptide synthetase (NRPS) and NRPS- polyketide synthase hybrid (NRPS/PKS) units, *clbK* and *clbJ* (Figure 7a, 7b). Interestingly, this fusion splits the pks island into two distinct and conserved majority-minority variants that are not congruent with the phylogeny of the pks+ isolates and do not appear to have originated from a clonal expansion (Figure 7b). The fusion is stably maintained in the population, however, being present in 14% of the pks+ *E. coli* (134 out of 954) with minor other changes in other pks island genes from isolates carrying the fused genes. This suggests that the *clbK-J* fusion may be under balancing selection, possibly with a different functional profile from the major variant, but such that both variants are competitive and selected for.

**Figure 7.**
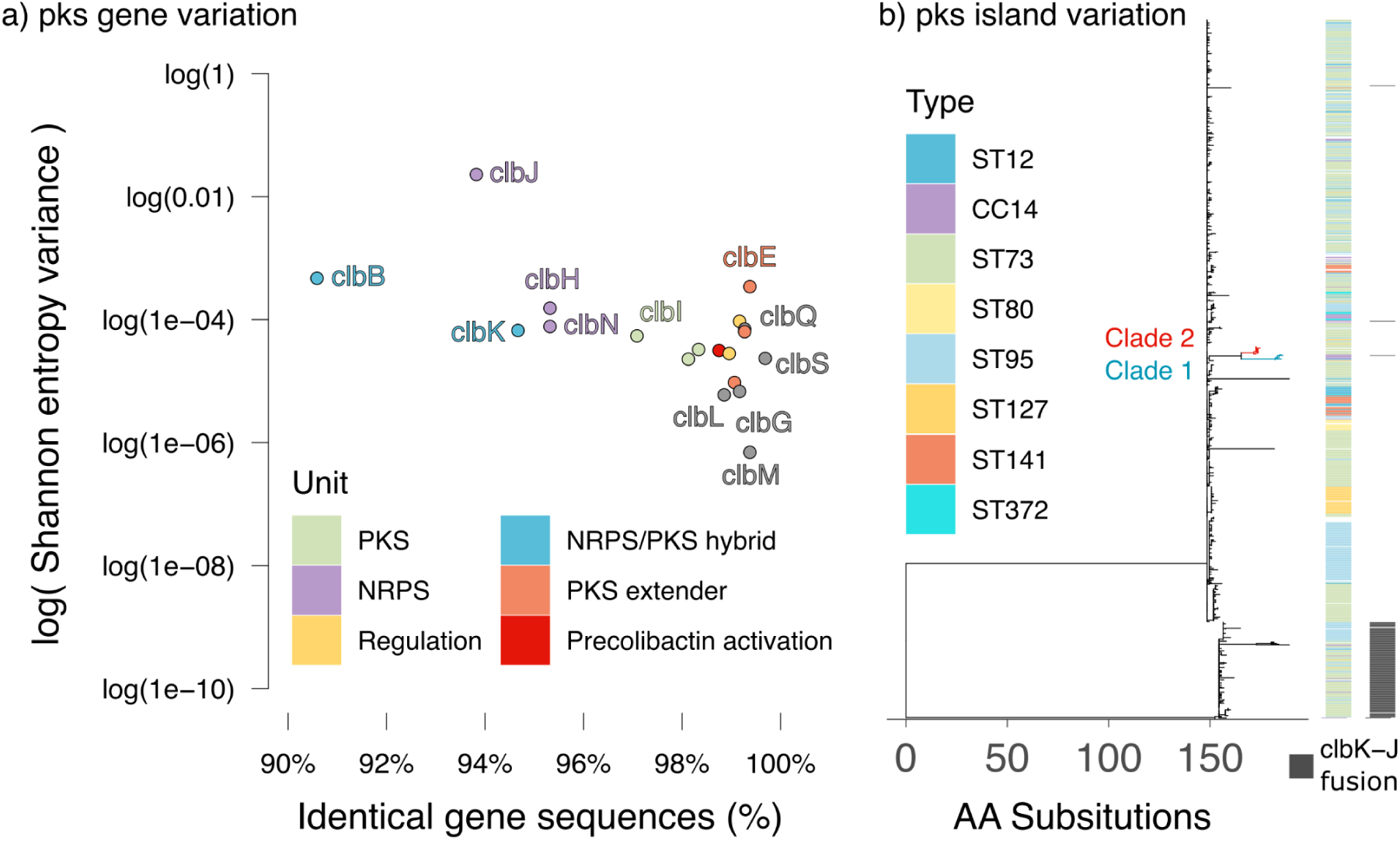
Amino acid sequence variation in the pks island. Panel a) shows the percentage of identical amino acid sequences and Shannon entropy variance (on a log-scale) for each gene in the pks island. The colours group the genes into units based on polyketide synthases (PKS; *clbC*, *clbI*, *clbO*), nonribosomal peptide synthases (NRPS; *clbH*, *clbJ*, *clbN*), NRPS/PKS hybrid (*clbB*, *clbK*), regulation (*clbR*, *clbA*), PKS extender (*clbD*, *clbE*, *clbF*), or activation (*clbP*). Genes not assigned to a unit are all labelled and coloured grey. Panel b) shows a phylogeny inferred on the concatenated amino acid sequences for all genes in the pks island. On the tree, the first metadata block shows the sequence type or clonal complex of the most common pks+ isolates. The second metadata block shows isolates with fused *clbK* and *clbJ* genes. The scale on the horizontal axis shows the branch length times the total number of sites in the concatenated amino acid sequences (16 976).

Outside of the *clbK-J* fusion, the pks island phylogeny shows a small expansion of two clades (“Clade 1” and “Clade 2” in Figure 7b and Supplementary Figure 2) which carry a number of conserved amino acid substitutions in 14 out of 19 genes that differentiate them from other pks+ *E. coli*. The two clades carry the same conserved versions of the main transcriptional activator *clbR* ^21^, the self-resistance gene *clbS* ^26,44^, the transporter *clbM* responsible for transporting precolibactin from the cytoplasm into the periplasm ^45^, an amidase *clbL* that dimerizes the precolibactin ^46^, and a freestanding acyltransferase *clbG* that transfers a PKS extender unit constructed by *clbD-F* to *clbK* ^47^. These same genes overall contain very few changes across the whole pks+ E. coli population (Figure 7a, Supplementary Figure 2) in our collection and are likely critical for the fitness of the bacteria producing the toxin.

## Discussion

Microbiota colonizing the human gut form a delicate ecological system whose stability is driven by competition and cooperation between its different transient and persisting members^48,49^. Bacteria in particular produce a multitude of toxins whose primary function is to displace other strains and species competing for the same resources, allowing the producing cells to establish dominance in a particular ecological context. Of these toxins, colibactin has recently received most attention due to its inadvertent genotoxic effect on eukaryotic DNA ^50–52^, as a byproduct of competition in the gut microbiome.

The general use of antibiotics since the introduction of penicillins has undoubtedly exerted an increasing selective pressure on the bacteria frequently colonising humans to either become resistant or decline in prevalence. Against this background, the dominant colibactin-producing *E. coli* stand in stark contrast to the trend, generally presenting with a markedly more susceptible phenotype but nevertheless thriving as both commensals and as leading causes of urinary tract and bloodstream infections in high-income settings ^10,14,15^. By combining recent insights from the *E. coli* plasmidome analysis ^16^, our study sheds further light into this contradiction, showing that instead of tolerating a wider class of antimicrobials, these bacteria have adapted to displacing competitors through the production of an arsenal of toxins including colibactin and various colicins. Similarly to the latter which often move around with plasmids, the pks island appears to move around via horizontal gene transfer from an integrative conjugative element that is present in at least *K. pneumoniae* ^43,53^, although in this species the pks island does not appear to confer a significant fitness boost as demonstrated by the absence of colibactin genes in samples from a One Health study conducted in Italy ^54^.

Combined, our phylogenetic analyses support the conclusion that the pks island was already acquired by the most recent common ancestor of all pks+ *E. coli* in the B2 phylogroup. Noting that pks+ B2 strains are the dominant cause of urinary tract and bloodstream infections among all *E. coli* , the evidence points to a hypothesis of their superiority in asymptomatic gut colonisation and presence in the bladder (bacteriuria) against other *E. coli* under lack of pressure from modern classes of broad-spectrum antibiotics, such as 3rd generation cephalosporins. When selection pressure from modern antibiotics is present, the fitness of the antibiotic susceptible pks+ *E. coli* decreases, opening the door for the widely established MDR strains that do not carry the pks island and have acquired resistance to colibactin around the same time they acquired resistance to the critical antibiotic classes. In the MDR strains, the additional fitness cost of colibactin resistance is then either negated by selection pressure from antimicrobial usage on susceptible strains or overcome by the competitive advantage over the pks+ *E. coli* that remain in circulation in parts of the population. An exception to this is ST1193, which according to previous studies emerged in South East Asia and then started expanding globally a decade ago ^36^. The lack of competition from the mainly susceptible colibactin-producers in this region where the use of in particular beta-lactam antibiotics is much higher than in Europe and North America, would have facilitated establishment of ST1193 without colibactin resistance.

Contrasting *Klebsiella,* we found evidence supporting that in *E. coli* co-evolution between colibactin production and resistance in clinically dominant lineages is a critical factor for becoming endemically circulating at high prevalence. This is apparent in the MDR lineages ST131 and ST69, which have acquired the *clbS*-like gene via a plasmid carrying a *clbS*-like transposon before undergoing clonal expansion reported in previous studies ^10,14,16^ and in ST131 C1/C2 the resistance gene got further integrated to the chromosome. It is reasonable to assume that without colibactin resistance, the constant competitive pressure from established colibactin-producing lineages would otherwise prevent the MDR lineages from establishing themselves in the host community under ecological settings where antibiotic usage pressure is low. The presence of *clbS-*like genes in a wide range of bacteria and structural similarity to the colibactin self-resistance protein *clbS* is recognized in the literature ^23,55^, but existing studies have not investigated the distribution of these genes in bacterial strains that are the main competitors of colibactin producers. Interestingly, a ClbS-like protein is also reported to induce cell death in neighbouring colonies when produced by *Paenibacillus dendritiformis* in dense colonies ^56^, and as such may have a role in *E. coli* that goes beyond protection from colibactin if this observation generalizes to other bacterial species.

In combination with the strong association of *clbS*-like genes with pks- *E. coli* in regions with widespread pks+ *E. coli*, the structural similarity is highly suggestive of this protein granting a degree of resistance to colibactin. Additionally, since the *clbS*-like genes can be transmitted via a plasmid and were acquired by multiple MDR clades relatively recently in their evolutionary history, the presence of this gene may constitute a key component in success of other clinically relevant bacterial pathogens that have expanded clonally in high-income countries in the antibiotic era. The *clbS*-like genes are a highly probable fitness determinant in bacteria that have to defend themselves against colibactin and may even provide a more effective defence mechanism than the *clbS* gene itself, in particular ecological settings. However, future experimental studies into *clbS*-like genes would be needed to conclusively establish its role and significance in pathogenic bacteria.

The pks island genes in *E. coli* are remarkably conserved ^33,57^ and we found this especially holds for genes that are responsible for activating the precolibactin or exporting it out of the cytoplasm. Changes to these genes are likely purged from the population, or are lethal to the producing cells, making them excellent targets for future studies that require stable target sequences, such as designing PCR primers to screen for colibactin production without genomic sequencing. Outside of the conserved genes, earlier studies have reported that variation in the pks island genes is limited to the *clbK* and *clbJ* fusion and that this fusion prevents the genotoxic activity of colibactin ^33^. Our results confirm that the *clbK-J* fusion is the only large-scale change in a collection that adequately represents the circulating population of *E. coli*. However, the remarkable conservation within the pks island in isolates with the fused genes strongly implies that this variant of the toxin is still functional against at least other bacteria. Additionally, the fusion does not appear to have resulted from clonal expansion or as an acquisition from a different source, but rather has evolved under balancing selection that stabilizes the minority variant in the population. Considering that the NRPS/PKS hybrid produced by *clbK* incorporates the α-aminoketone unit responsible for sequence specificity of the DNA crosslinking by colibactin ^46,58^, modifications to *clbK* and the other genes involved in NRPS and PKS production may be advantageous for pks+ *E. coli* in circumventing resistance conferred by ClbS-like proteins in non-producing bacteria, which is another promising target for further experimental research.

In this study, we have shown that the evolution of colibactin production in *E. coli* is tightly intertwined with the spread of antimicrobial resistant non-producing lineages and that the genotypes of lineages with the pks island of genes are highly adapted to producing the toxin at the cost of lacking the ability to acquire multi-drug resistance phenotypes. In parallel, the introduction of colibactin resistance genes in multi-drug resistant non-producing *E. coli* is clearly linked with their expansion in the recent decades. Our findings have significant implications on understanding the degrees of freedom bacteria are facing when evolving strategies in order to establish endemicity under different ecological settings.

## Methods

### Hybrid short- and long-read assemblies from 1 991 *E. coli* isolated from bloodstream infections

We used 1 991 *E. coli* hybrid Illumina+Nanopore assemblies from a previous study ^16^ and available from the European Nucleotide Archive under accession number <u>PRJEB45354</u> and <u>PRJEB57633</u>. Briefly, the isolates in this study were collected as part of the Norwegian Surveillance System for Resistant Microbes (NORM) study on extraintestinal pathogenic *E. coli* ^10^ and selected for Nanopore sequencing using an approach that ensured the chosen isolates adequately represent the accessory genome diversity in the NORM population ^16^. This selection was performed based on Illumina sequencing data obtained from 3 254 isolates in an earlier study ^10^. Additionally, all isolates belonging to the major extraintestinal pathogenic *E. coli* lineages ST69, ST73, ST95, and ST131 were selected for Nanopore sequencing and subsequent assembly. Hybrid assembly was performed using a custom pipeline (https://gitlab.com/sirarredondo/hybrid_assembly) built around Unicycler v0.4.7 ^59^.

### Genome annotation

We re-annotated the hybrid assembly sequences using bakta v1.10.4 ^60^ against the full database (v5.1, dated 2024-01-19). Contigs shorter than 200 base pairs were excluded. The annotation files generated in our study are available from Zenodo under doi: 10.5281/zenodo.17453317.

### Identifying ClbS-like proteins based on nucleotide and structural similarity

We extracted a total of 125 465 protein structure predictions from the NORM hybrid assemblies by creating a list of all unique UniRef 100 identifiers as assigned by bakta and querying the AlphaFold application programming interface hosted at the European Bioinformatics Institute ^61^. The resulting structure predictions were aligned against the structure of the colibactin self protection protein ClbS (AF-Q0P7K8-F1-model_v4) with foldmason release 3-954d202 ^62^ to obtain the mean local difference distance test (mLDDT) value between ClbS and all predicted protein structures. Based on the mLDDTs, we designated all proteins with a mLDDT higher than 80% as a *clbS*-like gene. This threshold was chosen because of an observed gap in the distribution of the mLDDT values (available under doi: 10.5281/zenodo.17017465) and a mismatch in the structural prediction for the protein with the next highest mLDDT value of 63% (Supplementary Figure 3).

### Determining presence and absence of pks island genes and the *clbS*-like gene

Presence or absence of the pks island was determined by counting the number of genes from the pks island (*clbA* to *clbS*) present in the bakta annotations for each hybrid assembly. An assembly was considered to carry the pks island if it produced at least 18 hits to the 19 genes, with the value 18 chosen because of the presence of the *clbK* and *clbJ* fusion in some assemblies. Two assemblies contained an intermediate number of pks island genes (2 out of 19) and were discarded from further analyses (GCA_964045745.1 carried *clbS* and *clbQ*, and GCA_964048735.1 carried *clbK* and *clbJ*). Assemblies with 0 pks island genes were designated pks negative.

The presence of *clbS*-like genes was first determined by searching for the presence of any of the gene sequences found using the structural search described earlier in the annotation files for each assembly. Then, we extracted the nucleotide sequences of each *clbS* and *clbS*-like gene and dereplicated them based on the UniRef 100 identifier and used the find subcommand in kbo-cli v0.2.1 with default parameters ^63^ to screen for possible variants without an UniRef identifier and at least 90% coverage and 99% identity to any of the *clbS* or *clbS*-like sequences. The *clbS* and *clbS*-like sequences identified in our study are available under doi: https://doi.org/10.5281/zenodo.17017465. Metadata showing presence and absence of the pks and clbS-like genes in each isolate is available in Supplementary Table 1.

### Assigning pks and *clbS*-like genes to chromosome, plasmid, or prophage sequences

We used genomad v1.11.1 ^64^ to compute the probability of each contig in the hybrid assemblies belonging to a plasmid or virus sequence and to determine the presence of prophage sequences in them. A contig was considered a plasmid or virus sequence if it had a score of at least 0.90 from genomad. No contig containing the pks, *clbS*, or *clbS*-like genes was assigned to a non-integrated bacteriophage sequence. Prophage regions were determined using the same threshold for the score. For contigs that were predicted as plasmid sequences, we additionally extracted the plasmid type information from an earlier study ^16^ by merging the annotations. Contig type predictions with presence and absence of at least one *clbS*-like gene are provided in Supplementary Table 2.

### Plasmid typing

Plasmid types were obtained from a previously published study ^16^. Briefly, the set of circularised plasmid sequences was deduplicated using cd-hit-est v4.8.1^65,66^ with sequence identity threshold 99%, alignment coverage 90%, and length difference cutoff at 90%. This yielded 2560 plasmid sequence clusters and one representative plasmid sequence was chosen from each. The set of representative sequences was used as input to mge-cluster v1.1.0 ^67^ and run with varying parameters. The results from runs with different parameters were merged into a co-occurrence network by thresholding the probability of a representative sequence belonging to a cluster in each run with at least 80% probability. The co-occurrence network was converted into community assignments by using the Louvain algorithm ^68^ from the R igraph ^69^ package to identify dense communities with a modularity threshold of 20%. These results were used to define the two digit plasmid type with the first digit indicating the graph component (non-singleton graph without connections to outside of it) and the second digit indicating the community assignment within the component identified by the Louvain algorithm. Redundant plasmid sequences removed by cd-hit-est in the first step were assigned to the same plasmid type as their representative sequence.

Finally, non-circularised plasmid sequences were assigned to a plasmid type by using the --existing option from mge-cluster to predict the coordinates for the non-circularised sequences in the 2D embedding defined by the circularised plasmids, and assigning the non-circularised sequences to the closest (Euclidean distance) plasmid type.

### Pangenome analysis

Pangenome analysis for the NORM hybrid assemblies and core gene alignment was performed using panaroo v1.5.2 ^70^ with the “--clean-mode strict” and “--remove-invalid-genes” options. Panaroo was run separately for 1) all 1 991 hybrid assemblies, 2) all pks+ assemblies, and 3) all pks- assemblies. The output files are available from Zenodo under doi: 10.5281/zenodo.17521041.

### Synteny analysis for the pks island neighbourhood

The pks island neighbourhood was defined as the set of genes between *cobU* and *mtfA*. This region was extracted from nine representative hybrid assembly annotations, chosen randomly from the assemblies with a fully intact set of the 19 pks island genes and fed to clinker v0.0.31 to perform the synteny analysis ^71^. The files used in performing the clinker analysis are available under doi: 10.5281/zenodo.17629452.

### Species-wide phylogeny

The species-wide phylogeny in Figure 2 was generated from the pan-genome core alignment inferred with panaroo v1.5.2 using VeryFastTree v4.0.0 ^72,73^ using a Gamma+GTR+CAT model with 20 categories, no bootstrap support values, the -fastest option, and 32-bit floating point precision. The phylogeny was visualized with ggplot2 and ggtree ^74^. The pan-genome core alignment used to build the phylogeny and the output files are available under doi: 10.5281/zenodo.17608175.

### Phylogenetic dating of the major *E. coli* lineages

Dated phylogenies for ST69, ST73, ST95, and ST131 were obtained from previous studies ^10,16^. A dated phylogeny was not available for the CC14 isolates, and was generated using the following approach: the sequencing reads from each CC14 isolate were aligned against the *E. coli* M1/5 sequence belonging to ST550 in CC14 (GCF 013390265.1) with bwa mem v0.7.18^75^. The alignments were sorted and indexed with samtools v1.21 ^76^ and fed to freebayes v1.3.8 ^77^ to infer variants between the reference and each isolate. The freebayes results were filtered with bcftools v1.22 ^78^ for results with quality higher or equal to 100 and converted to a reference based alignment using the consensus subcommand from bcftools. Then, the reference based alignments were concatenated and raxml-ng v1.2.2 with the GTR+G20 model, 100 random and 100 parsimony starting trees, minimum branch length 10^-12^ and safe branch length optimization ^79^ was used to infer a maximum likelihood tree from the alignment. This tree was then used as a starting tree for gubbins v3.4.3 with extensive search with 10 iterations and the GTRGAMMA model using iqtree 2.4.0 as the tree builder ^80,81^ to infer a recombination free phylogeny from a core genome alignment generated with snp-sites from the full reference based alignment ^82^. Finally, this tree was dated using the bactdating ^83^ R package (GitHub commit 8825600) with the additive uncorrelated relaxed clock model ^84^. The date of the most recent common ancestor was confirmed statistically significant at significance level 0.01 using a permutation test shuffling the isolation dates with 1 000 permutations. Dated phylogenies for ST69, ST74, ST95, and ST131 are available from the previous study at https://gitlab.com/sirarredondo/expec_plasmidome. The files used in constructing the dated phylogeny for CC14 are available under doi: 10.5281/zenodo.17629103.

### Variation in pks island genes

We extracted the amino acid sequences for each gene in the pks island (*clbA-clbS*) from the bakta annotations for each pks+ hybrid assembly and performed multiple sequence alignment separately on each gene using mafft v7.5.2 with the --globalpair and --maxiterate 1000 options ^85^. The per-gene alignments were concatenated in the order they typically appear in (*clbA, clbR, clbB*-*clbS*) and the concatenated file was used to infer the phylogeny presented in Figure 7 using raxml-ng with the LG+G8+F model from 100 random and 100 parsimony starting trees with minimum branch length 10^-12^ and safe branch length optimization ^79^. Per-gene trees were constructed by applying the same approach on the gene-wise multiple sequence alignments from mafft. The tree files and the input multiple sequence alignments are available under doi: 10.5281/zenodo.17629325.

## Data availability

Output files from the analyses presented are available from Zenodo and linked to under the relevant Methods section headers. A Microreact ^86^ project providing an interactive view of the species-wide phylogeny in Figure 2 is available from https://microreact.org/project/norm-colibactin-coevolution. Source files for all figures and supplementary figures and interactive views for Figures 5 and 6 are available under doi: 10.5281/zenodo.17607676

## Author contributions

TM was responsible for formal analysis and visualisations. TM and JKC performed the investigation into *clbS*-like gene carriage and transfer in plasmids. TM and SM analysed *clbS*-like gene carriage in prophage regions. TM and HAT investigated the mobile element origin for pks island genes from *Klebsiella*. RAG provided contextualization for the NORM collection. TM coordinated the project. TM and JC wrote the initial draft and conceptualised the study. JC acquired funding. All authors contributed to the writing, reviewing, and editing of the manuscript.

## Acknowledgements

T.M. and J.C. were partially funded by the Research Council of Norway, grant no. 299941 and by Wellcome Trust grant no. 220540/Z/20/A to the Wellcome Sanger Institute.

The authors wish to acknowledge CSC – IT Center for Science, Finland, for computational resources.

## References

1. Pleguezuelos-Manzano, C. et al. Mutational signature in colorectal cancer caused by genotoxic pks *E. coli*. Nature 580, 269–273 (2020).

2. Dziubańska-Kusibab, P. J. et al. Colibactin DNA-damage signature indicates mutational impact in colorectal cancer. Nature Medicine 26, 1063–1069 (2020).

3. Lee-Six, H. et al. The landscape of somatic mutation in normal colorectal epithelial cells. Nature 574, 532–537 (2019).

4. Mäklin, T. et al. Geographical variation in the incidence of colorectal cancer and urinary tract cancer is associated with population exposure to colibactin-producing *Escherichia coli*. Lancet Microbe 6, 101015 (2025).

5. Díaz-Gay, M. et al. Geographic and age variations in mutational processes in colorectal cancer. Nature 643, 230–240 (2025).

6. Shrestha, E. et al. Oncogenic gene fusions in nonneoplastic precursors as evidence that bacterial infection can initiate prostate cancer. Proceedings of the National Academy of Sciences of the United States of America 118, (2021).

7. Agrawal, R. et al. Colibactin exerts androgen-dependent and -independent effects on prostate cancer. European Urology Oncology 8, 716–730 (2025).

8. Chagneau, C. V. et al. Uropathogenic *E. coli* induces DNA damage in the bladder. PLOS Pathogens 17, e1009310 (2021).

9. Li, D. et al. Dominance of sequence types ST73, ST95, ST127 and ST131 in Australian urine isolates: a genomic analysis of antimicrobial resistance and virulence linked to F plasmids. Microbial Genomics 9, (2023).

10. Gladstone, R. A. et al. Emergence and dissemination of antimicrobial resistance in *Escherichia coli* causing bloodstream infections in Norway in 2002-17: a nationwide, longitudinal, microbial population genomic study. Lancet Microbe 2, e331–e341 (2021).

11. Chagneau, C. V. et al. The pks island: a bacterial Swiss army knife? Colibactin: beyond DNA damage and cancer. Trends in Microbiology 30, 1146–1159 (2022).

12. Nhu, N. T. K. et al. High-risk clones that cause neonatal meningitis and association with recrudescent infection. eLife 12, (2024).

13. Kallonen, T. et al. Systematic longitudinal survey of invasive *Escherichia coli* in England demonstrates a stable population structure only transiently disturbed by the emergence of ST131. Genome Research 27, 1437–1449 (2017).

14. Pöntinen, A. K. et al. Modulation of multidrug-resistant clone success in *Escherichia coli* populations: a longitudinal, multi-country, genomic and antibiotic usage cohort study. Lancet Microbe 5, e142–e150 (2024).

15. Mäklin, T. et al. Strong pathogen competition in neonatal gut colonisation. Nature Communications 13, 7417 (2022).

16. Arredondo-Alonso, S. et al. Plasmid-driven strategies for clone success in *Escherichia coli*. Nature Communications 16, 2921 (2025).

17. Palomino, A. et al. Metabolic genes on conjugative plasmids are highly prevalent in *Escherichia coli* and can protect against antibiotic treatment. The ISME Journal 17, 151–162 (2023).

18. Connor, C. H. et al. Multidrug-resistant *E. coli* encoding high genetic diversity in carbohydrate metabolism genes displace commensal *E. coli* from the intestinal tract. PLOS Biology 21, e3002329 (2023).

19. Addington, E., Sandalli, S. & Roe, A. J. Current understandings of colibactin regulation. Microbiology 170, (2024).

20. Tronnet, S. et al. Iron homeostasis regulates the genotoxicity of *Escherichia coli* that produces colibactin. Infection and Immunity 84, 3358–3368 (2016).

21. Wallenstein, A., et al. ClbR Is the key transcriptional activator of colibactin gene expression in *Escherichia coli*. mSphere 5, (2020).

22. Bossuet, N. et al. Oxygen concentration modulates colibactin production. Gut Microbes 15, 2222437 (2023).

23. Silpe, J. E., Wong, J. W. H., Owen, S. V., Baym, M. & Balskus, E. P. The bacterial toxin colibactin triggers prophage induction. Nature 603, 315–320 (2022).

24. Wong, J. J., et al. *Escherichia coli* BarA-UvrY regulates the pks island and kills *Staphylococci* via the genotoxin colibactin during interspecies competition. PLOS Pathogens 18, e1010766 (2022).

25. Tronnet, S., et al. The genotoxin colibactin shapes gut microbiota in mice. mSphere 5, (2020).

26. Bossuet-Greif, N., et al. *Escherichia coli* ClbS is a colibactin resistance protein. Molecular Microbiology 99, 897–908 (2016).

27. Hancock, V., Dahl, M. & Klemm, P. Probiotic *Escherichia coli* strain Nissle 1917 outcompetes intestinal pathogens during biofilm formation. Journal of Medical Microbiology 59, 392–399 (2010).

28. Maltby, R., Leatham-Jensen, M. P., Gibson, T., Cohen, P. S. & Conway, T. Nutritional basis for colonization resistance by human commensal *Escherichia coli* strains HS and Nissle 1917 against *E. coli* O157:H7 in the mouse intestine. PLOS One 8, e53957 (2013).

29. Hare, P. J., Englander, H. E. & Mok, W. W. K. Probiotic *Escherichia coli* Nissle 1917 inhibits bacterial persisters that survive fluoroquinolone treatment. Journal of Applied Microbiology 132, 4020–4032 (2022).

30. Wernke, K. M. et al. Structure and bioactivity of colibactin. Bioorganic & Medicinal Chemistry Letters 30, 127280 (2020).

31. Levy, S. et al. Colibactin genes are highly prevalent in the developing infant gut microbiome. medRxiv (2025) doi:10.1101/2025.08.12.25333511.

32. Tsunematsu, Y. et al. Mother-to-infant transmission of the carcinogenic colibactin-producing bacteria. BMC Microbiology 21, 235 (2021).

33. Auvray, F. et al. Insights into the acquisition of the pks island and production of colibactin in the *Escherichia coli* population. Microbial Genomics 7, (2021).

34. Be’er, A. et al. Lethal protein produced in response to competition between sibling bacterial colonies. Proceedings of the National Academy of Sciences 107, 6258–6263 (2010).

35. Manges, A. R. et al. Global extraintestinal pathogenic *Escherichia coli* (ExPEC) lineages. Clinical Microbiology Reviews 32, (2019).

36. Pitout, J. D. D., Peirano, G., Chen, L., DeVinney, R. & Matsumura, Y. *Escherichia coli* ST1193: following in the footsteps of *E. coli* ST131. Antimicrobial Agents and Chemotherapy 66, e0051122 (2022).

37. Koh, E.-I., Robinson, A. E., Bandara, N., Rogers, B. E. & Henderson, J. P. Copper import in Escherichia coli by the yersiniabactin metallophore system. Nature Chemical Biology 13, 1016–1021 (2017).

38. Katumba, G. L., Tran, H. & Henderson, J. P. The Yersinia High-Pathogenicity Island Encodes a Siderophore-Dependent Copper Response System in Uropathogenic Escherichia coli. mBio (2022) doi:10.1128/mBio.02391-21.

39. Heffernan, J. R. et al. Yersiniabactin is a quorum-sensing autoinducer and siderophore in uropathogenic Escherichia coli. mBio (2024) doi:10.1128/mbio.00277-23.

40. Lawrence, J. G. & Roth, J. R. The cobalamin (coenzyme B12) biosynthetic genes of Escherichia coli. Journal of Bacteriology (1995) doi:10.1128/jb.177.22.6371-6380.1995.

41. Escalante-Semerena, J. C. & Warren, M. J. Biosynthesis and Use of Cobalamin (B12). EcoSal Plus (2008) doi:10.1128/ecosalplus.3.6.3.8.

42. Wookey, P. & Rosenberg, H. Involvement of inner and outer membrane components in the transport of iron and in colicin B action in Escherichia coli. Journal of Bacteriology 133, 661–666 (1978).

43. Lam, M. M. C. et al. Genetic diversity, mobilisation and spread of the yersiniabactin-encoding mobile element ICEKp in *Klebsiella pneumoniae* populations. Microbial Genomics 4, (2018).

44. Tripathi, P. et al. ClbS Is a cyclopropane hydrolase that confers colibactin resistance. Journal of the American Chemical Society (2017) doi:10.1021/jacs.7b09971.

45. Mousa, J. J. et al. MATE transport of the *E. coli*-derived genotoxin colibactin. Nature Microbiology 1, 1–7 (2016).

46. Jiang, Y. et al. Reactivity of an unusual amidase may explain colibactin’s DNA cross-linking activity. Journal of the American Chemical Society (2019) doi:10.1021/jacs.9b02453.

47. Zha, L., Wilson, M. R., Brotherton, C. A. & Balskus, E. P. Characterization of polyketide synthase machinery from the pks island facilitates isolation of a candidate precolibactin. ACS Chemical Biology (2016) doi:10.1021/acschembio.6b00014.

48. Spragge, F. et al. Microbiome diversity protects against pathogens by nutrient blocking. Science 382, eadj3502 (2023).

49. Wilde, J., Slack, E. & Foster, K. R. Host control of the microbiome: mechanisms, evolution, and disease. Science 385, eadi3338 (2024).

50. Nougayrède, J.-P., et al. *Escherichia coli* induces DNA double-strand breaks in eukaryotic cells. Science 313, 848–851 (2006).

51. Cuevas-Ramos, G., et al. *Escherichia coli* induces DNA damage in vivo and triggers genomic instability in mammalian cells. Proceedings of the National Academy of Sciences of the United States of America 107, 11537–11542 (2010).

52. Vizcaino, M. I. & Crawford, J. M. The colibactin warhead crosslinks DNA. Nature Chemistry 7, 411–417 (2015).

53. Wami, H. et al. Insights into evolution and coexistence of the colibactin- and yersiniabactin secondary metabolite determinants in enterobacterial populations. Microbial Genomics 7, (2021).

54. Thorpe, H. A. et al. A large-scale genomic snapshot of *Klebsiella* spp. isolates in Northern Italy reveals limited transmission between clinical and non-clinical settings. Nature Microbiology 7, 2054–2067 (2022).

55. Tripathi, P. & Bruner, S. D. Structural basis for the interactions of the colibactin resistance gene product ClbS with DNA. Biochemistry 60, 1619–1625 (2021).

56. Taylor, J. D., Taylor, G., Hare, S. A. & Matthews, S. J. Structures of the DfsB protein family suggest a cationic, helical sibling lethal factor peptide. Journal of Molecular Biology 428, 554–560 (2016).

57. Suresh, A. et al. Evolutionary dynamics based on comparative genomics of pathogenic *Escherichia coli* lineages harboring polyketide synthase (pks) island. mBio 12, (2021).

58. Carlson, E. S., et al. The specificity and structure of DNA crosslinking by the gut bacterial genotoxin colibactin. bioRxiv (2025) doi:10.1101/2025.05.26.655968.

59. Wick, R. R., Judd, L. M., Gorrie, C. L. & Holt, K. E. Unicycler: Resolving bacterial genome assemblies from short and long sequencing reads. PLOS Computational Biology 13, e1005595 (2017).

60. Schwengers, O. et al. Bakta: rapid and standardized annotation of bacterial genomes via alignment-free sequence identification: Find out more about Bakta, the motivation, challenges and applications, here. Microbial Genomics 7, 000685 (2021).

61. Jumper, J. et al. Highly accurate protein structure prediction with AlphaFold. Nature 596, 583–589 (2021).

62. Gilchrist, C. L. M., Mirdita, M. & Steinegger, M. Multiple protein structure alignment at scale with FoldMason. bioRxiv (2024) doi:10.1101/2024.08.01.606130.

63. Mäklin, T., Alanko, J. N., Biagi, E. & Puglisi, S. J. Sequence alignment with *k*-bounded matching statistics. bioRxiv (2025) doi:10.1101/2025.05.19.654936.

64. Camargo, A. P. et al. Identification of mobile genetic elements with geNomad. Nature Biotechnology 42, 1303–1312 (2024).

65. Fu, L., Niu, B., Zhu, Z., Wu, S. & Li, W. CD-HIT: accelerated for clustering the next-generation sequencing data. Bioinformatics 28, 3150–3152 (2012).

66. Li, W. & Godzik, A. Cd-hit: a fast program for clustering and comparing large sets of protein or nucleotide sequences. Bioinformatics 22, 1658–1659 (2006).

67. Arredondo-Alonso, S., et al. Mge-cluster: a reference-free approach for typing bacterial plasmids. NAR Genomics and Bioinformatics 5, lqad066 (2023).

68. Blondel, V. D., Guillaume, J.-L., Lambiotte, R. & Lefebvre, E. Fast unfolding of communities in large networks. arXiv (2008) doi:10.48550/ARXIV.0803.0476.

69. Antonov, M., et al. graph enables fast and robust network analysis across programming languages. arXiv (2023) doi:10.48550/ARXIV.2311.10260.

70. Tonkin-Hill, G. et al. Producing polished prokaryotic pangenomes with the Panaroo pipeline. Genome Biology 21, 180 (2020).

71. Gilchrist, C. L. M. & Chooi, Y.-H. clinker & clustermap.js: automatic generation of gene cluster comparison figures. Bioinformatics 37, 2473–2475 (2021).

72. Piñeiro, C. & Pichel, J. C. Efficient phylogenetic tree inference for massive taxonomic datasets: harnessing the power of a server to analyze 1 million taxa. GigaScience 13, (2024).

73. Piñeiro, C., Abuín, J. M. & Pichel, J. C. Very Fast Tree: speeding up the estimation of phylogenies for large alignments through parallelization and vectorization strategies. Bioinformatics 36, 4658–4659 (2020).

74. Yu, G., Lam, T. T.-Y., Zhu, H. & Guan, Y. Two methods for mapping and visualizing associated data on phylogeny using ggtree. Molecular Biology and Evolution 35, 3041–3043 (2018).

75. Li, H. Aligning sequence reads, clone sequences and assembly contigs with BWA-MEM. arXiv (2013) doi:10.48550/ARXIV.1303.3997.

76. Li, H. et al. The sequence alignment/map format and SAMtools. Bioinformatics 25, 2078–2079 (2009).

77. Garrison, E. & Marth, G. Haplotype-based variant detection from short-read sequencing. arXiv (2012) doi:10.48550/ARXIV.1207.3907.

78. Danecek, P. et al. Twelve years of SAMtools and BCFtools. GigaScience 10, (2021).

79. Kozlov, A. M., Darriba, D., Flouri, T., Morel, B. & Stamatakis, A. RAxML-NG: a fast, scalable and user-friendly tool for maximum likelihood phylogenetic inference. Bioinformatics 35, 4453–4455 (2019).

80. Croucher, N. J. et al. Rapid phylogenetic analysis of large samples of recombinant bacterial whole genome sequences using Gubbins. Nucleic Acids Research 43, e15 (2015).

81. Minh, B. Q. et al. IQ-TREE 2: New models and efficient methods for phylogenetic inference in the genomic era. Molecular Biology and Evolution 37, 1530–1534 (2020).

82. Page, A. J. et al. SNP-sites: rapid efficient extraction of SNPs from multi-FASTA alignments. Microbial Genomics 2, e000056 (2016).

83. Didelot, X., Croucher, N. J., Bentley, S. D., Harris, S. R. & Wilson, D. J. Bayesian inference of ancestral dates on bacterial phylogenetic trees. Nucleic Acids Research 46, e134 (2018).

84. Didelot, X., Siveroni, I. & Volz, E. M. Additive Uncorrelated Relaxed Clock Models for the Dating of Genomic Epidemiology Phylogenies. Molecular Biology and Evolution 38, 307–317 (2021).

85. Katoh, K. & Standley, D. M. MAFFT multiple sequence alignment software version 7: improvements in performance and usability. Molecular Biology and Evolution 30, 772–780 (2013).

86. Argimón, S. et al. Microreact: visualizing and sharing data for genomic epidemiology and phylogeography. Microbial Genomics 2, e000093 (2016).

