## Supplementary figures and images for "Co-evolution between colibactin production and resistance is linked to clonal expansions in *Escherichia coli*"

### Supplementary Figure 1

#### Synteny between clbS like gene carrying plasmids and clbS like chromosomal regions

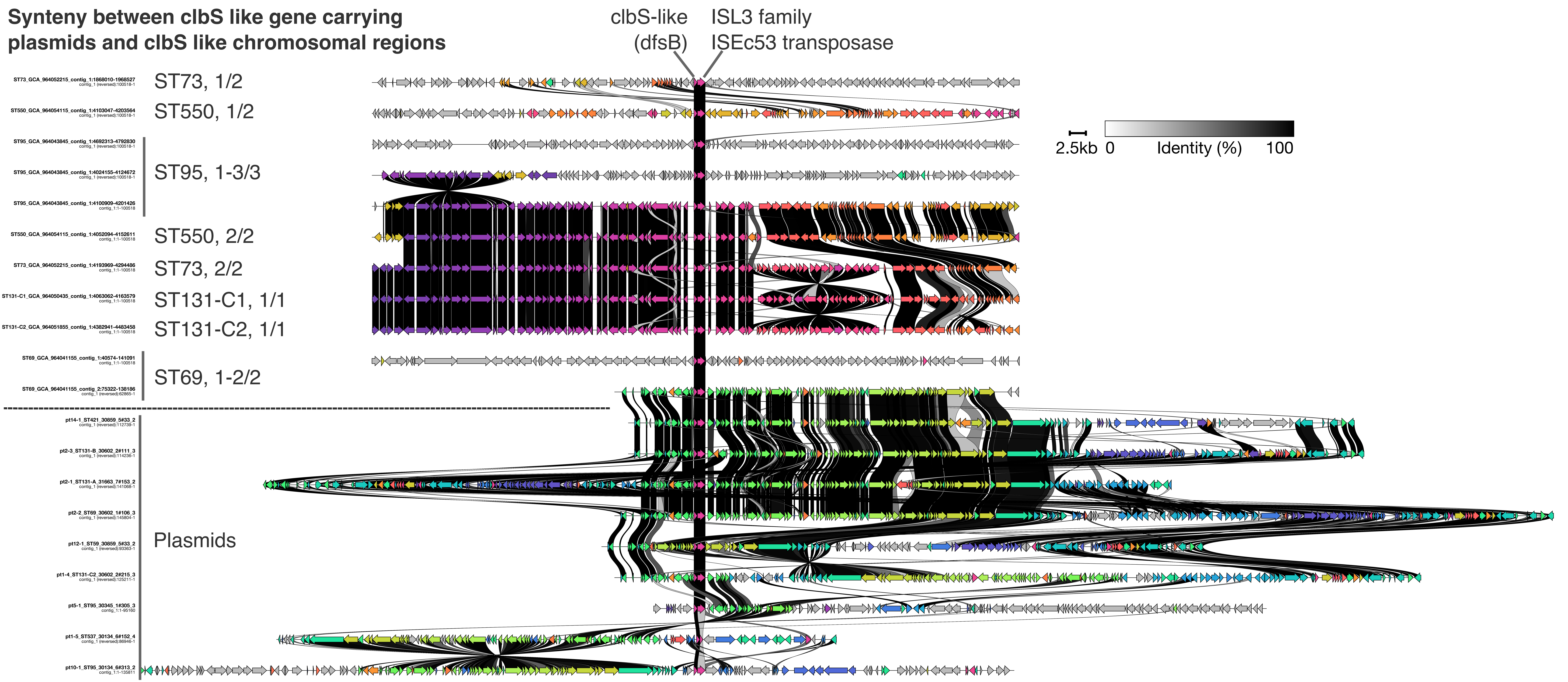

### Supplementary Figure 2

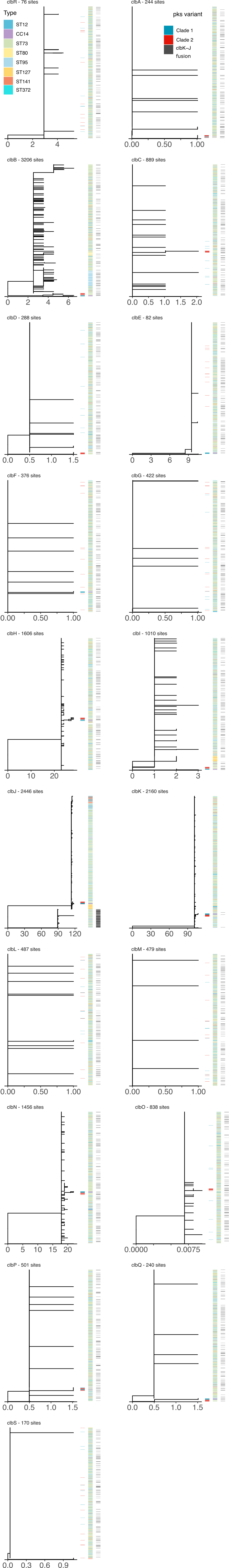

### Supplementary Figure 3

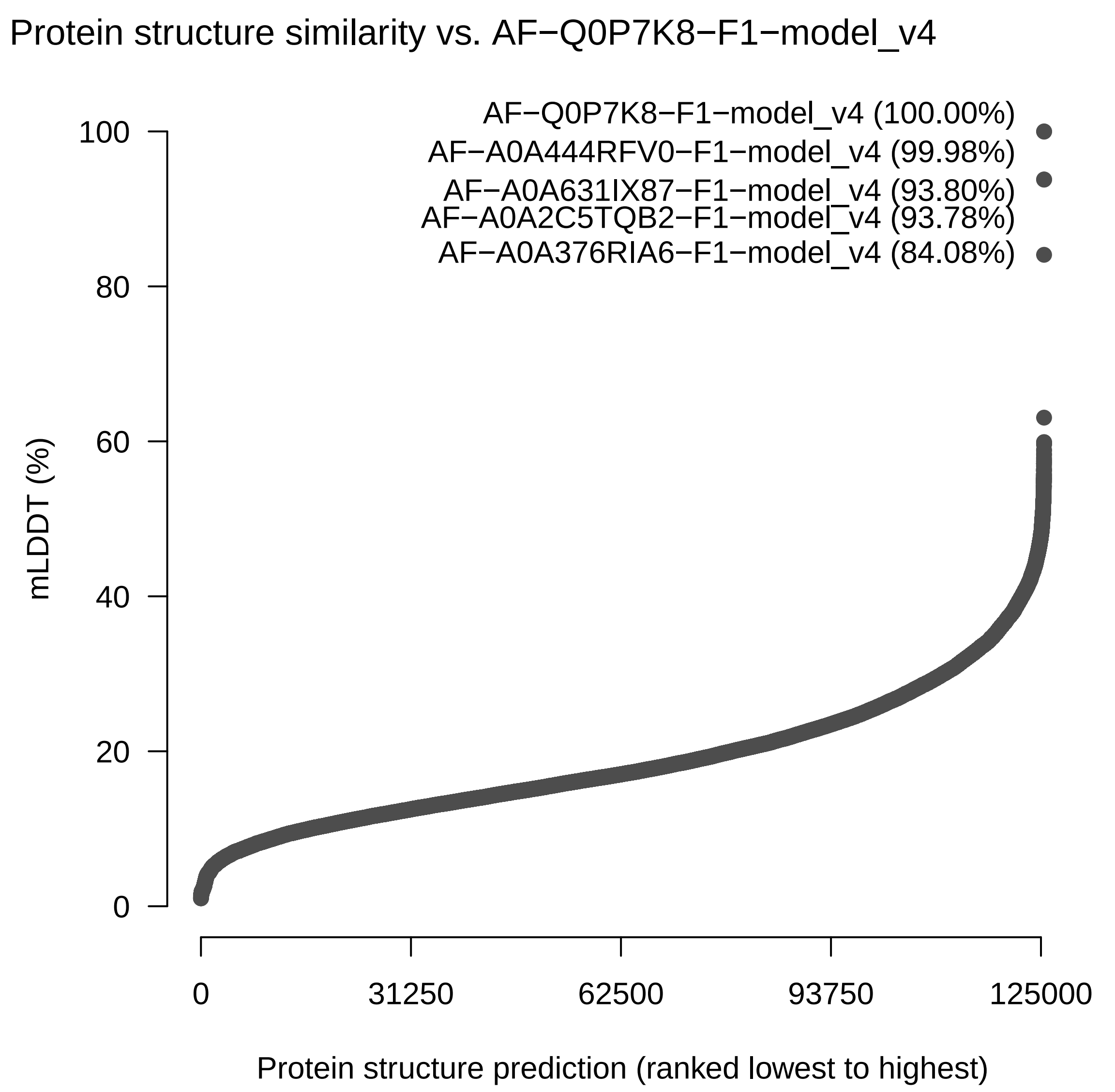
